# A mathematical framework to correct for compositionality in microbiome datasets

**DOI:** 10.1101/2025.05.29.656792

**Authors:** Samuel P. Forry, Stephanie L. Servetas, Jason G. Kralj, Monique E. Hunter, Jennifer N. Dootz, Scott A. Jackson

## Abstract

The increasing use of metagenomic sequencing (MGS) for microbiome analysis has significantly advanced our understanding of microbial communities and their roles in various biological processes, including human health, environmental cycling, and disease. However, the inherent compositionality of MGS data, where the relative abundance of each taxa depends on the abundance of all other taxa, complicates the measurement of individual taxa and the interpretation of microbiome data. Here we describe an experimental design that incorporates exogenous internal standards in routine MGS analyses to correct for compositional distortions. A mathematical framework was developed for using the observed internal standard relative abundance to calculate “Scaled Abundances” for native taxa that were (i) independent of sample composition and (ii) directly proportional to actual biological abundances. Through rigorous analysis of mock community and human gut microbiome samples, we demonstrate that Scaled Abundances outperformed traditional relative abundance measurements in both precision and accuracy and enabled reliable, quantitative comparisons of individual microbiome taxa across varied sample compositions and across a wide range of taxa abundances. By providing a pathway to accurate taxa quantification, this approach holds significant potential for advancing microbiome research, particularly in clinical and environmental health applications where precise microbial profiling is critical.

**Importance:** Metagenomic sequencing (MGS) analysis has become central to modern characterizations of microbiome samples. However, the inherent compositionality of these analyses often complicate interpretations of results. We present here an experimental design and corresponding mathematical framework that uses internal standards with routine MGS methods to correct for compositional distortions. We validate this approach for both amplicon and shotgun MGS analysis of mock communities and human gut microbiome (fecal) samples. By using internal standards to remove compositionality, we demonstrate significantly improved measurement accuracy and precision for quantification of taxa abundances. This approach is broadly applicable across a wide range of microbiome research applications.

## Introduction

Over the last several decades, the decreasing cost and increasing throughput of next generation sequencing measurements have made metagenomic sequencing (MGS) characterization a default strategy for microbiome analyses.^1–3^ Owing to the ubiquity of naturally occurring microbiomes throughout natural and man-made environments, MGS analyses have correlated microbiomes, and their resident microbes, with a variety of important phenomena, including human, animal, and plant health, renewable resources, infrastructure degradation, environmental biogeochemical cycling, and waste remediation, among others.^3–12^ Particularly in the field of health, many connections have been postulated between the human microbiome and various health and disease conditions such as obesity, gut health, autism, depression, or autoimmune disease.^4,6,8,13–18^

However, in spite of promising initial indications, many of the earliest correlations between microbiomes and health conditions have failed to bear fruit. For instance, questions have arisen about the relevance of the Firmicutes:Bacteroides ratio to obesity and gut health, about the existence of a ‘cancer microbiome’, and about the role of microbes in Autism.^19–22^ Even early in the Human Microbiome project, it was widely recognized that MGS results were much more reproducible (precise) than they were accurate (reflecting the underlying biology).^23,24^ Many comparisons have shown that analysis results vary significantly between samples and methodologies.^25–30^

One of the complications with drawing conclusions from MGS results lies in the data’s inherent compositionality.^25,27,30,31^ In short, the observed relative abundance of each taxa in a sample depends on both its actual abundance as well as the abundances of all other taxa in the sample. While aggregate compositional differences between samples can be accurately assessed, this severely limits the quantification of variations in the individual abundances of constituent taxa. Alternately, rigorous compositional data analysis strategies hold great promise for quantifying individual microbiome taxa abundances, correlating observed and actual abundances of constituent taxa, and allowing direct comparisons between microbiome samples with varied compositions.^32^

A variety of ratiometric analysis strategies have been applied in the context of microbiome MGS to try to correct for data compositionality.^32–35^ One effective strategy has been to use ratios of pairs of taxa relative abundances within each sample to correct for the effects of compositionality.^25,36^ Although this accurately accounts for compositionality, ratios of abundances of native taxa can be challenging to interpret and require that both taxa be present in all samples of interest. In wastewater biosurveillance for extracellular antimicrobial resistance genes by MGS, the routine addition of exogenous DNA as an internal standard has been demonstrated to improve MGS quantitation.^37,38^

In the current effort, we describe an experimental design for the routine inclusion of internal standards in microbiome MGS analyses. When genetic material from exogenous microbes is systematically added to samples, it serves as an internal reference standard to assess and mathematically correct for each sample’s composition. A rigorous framework for this analysis is provided along with several demonstrative examples. A previously reported systematic study of mock communities is re-analyzed here using the proposed internal standard framework. Then, a series of human gut microbiome samples are prepared and evaluated for two demonstration cases: (i) constant microbe concentrations with varied sample compositions; and (ii) systematically varied microbe concentrations with constant sample composition. In all cases, the results validate the proposed mathematical model for using internal standards to improve quantitative MGS analyses.

## Results and Discussion

### Mathematical framework

The analysis of microbiome samples using MGS varies among researchers but always requires a series of physical and bioinformatic manipulations, typically including sample acquisition, DNA extraction, library preparation, and next generation sequencing (NGS), as well as read trimming and taxonomic assignment.^1,25,29,30^ Importantly, each step contributes biases that accumulate throughout the measurement pipeline.^25,30^ For a locked-down method, these individual biases can be combined into a single aggregate bias term for each taxa that describes the analytical response through the entire protocol.^25^

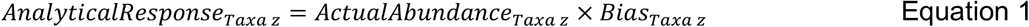

However, the analytical response is not directly observable from MGS results due to distortion arising from process saturation. Since NGS analyses arbitrarily limit the total number of DNA molecules that are analyzed, the measured relative abundance for each taxa depends on both the analytical response of that taxa (Eqn 1) as well as the aggregate responses of all taxa in the sample:

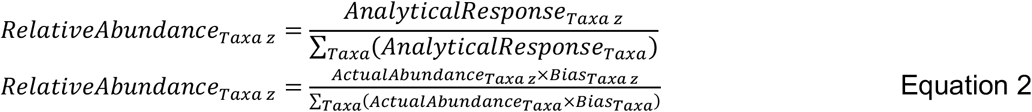

The denominator in Eqn 2 gives rise to the inherent compositional nature of MGS datasets since the measured relative abundance for each taxa is inextricably tied to relative abundance of every other taxa in the sample. So, for example, a measured increase in a taxa’s relative abundance could arise from either an increase in its actual biological abundance in the sample or from a decrease in the actual abundances of other Taxa. These two scenarios are phenomenologically distinct but generally indistinguishable experimentally, particularly as (i) the Actual Abundances and Biases remain unknown for most taxa in naturally occurring samples and (ii) biases vary substantially and unpredictably between individual taxa and various common protocols.^25,30^ Thus, while whole-sample comparisons based on measured relative abundances generally reflect aggregate compositional changes with some accuracy, at least within the context of locked-down protocols, the quantitative comparison of individual taxa relative abundances between samples is unreliable.

A variety of compositional data analysis approaches have been developed for or adapted to MGS datasets with some success.^25,32^ In general, these approaches recognize that the compositional denominator in Eqn 2 is identical for all taxa from a sample, and use ratios to remove or diminish the compositionality. For example, the ratio of the relative abundances of two native taxa from a sample will depend only on the actual taxa abundances and biases associated with those taxa and will be independent of the rest of the sample composition:^25^

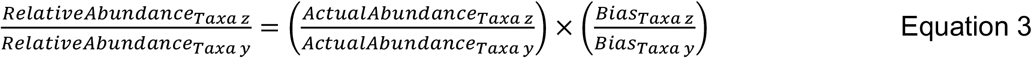

In the current effort, we propose an experimental design where an exogenous microorganism is systematically added into each sample to be analyzed as an internal standard (Figure 1). The strategy of employing internal standards to correct for analytical distortions between samples has been advantageously employed in such analyses as fluorescence quantitation, gene arrays, and metabolomic analyses.^39–41^ Basically, the internal standard is analyzed along with the rest of the sample throughout the entire measurement pipeline, and its measured abundance is similarly affected by sample-specific parameters such as compositionality. For application in MGS datasets, the internal standard can be readily distinguished from other taxa based on its unique genome sequence, and its observed relative abundance and known actual abundance (as specified by a protocol for its systematic addition) help correct for compositionality.

**Figure 1.**
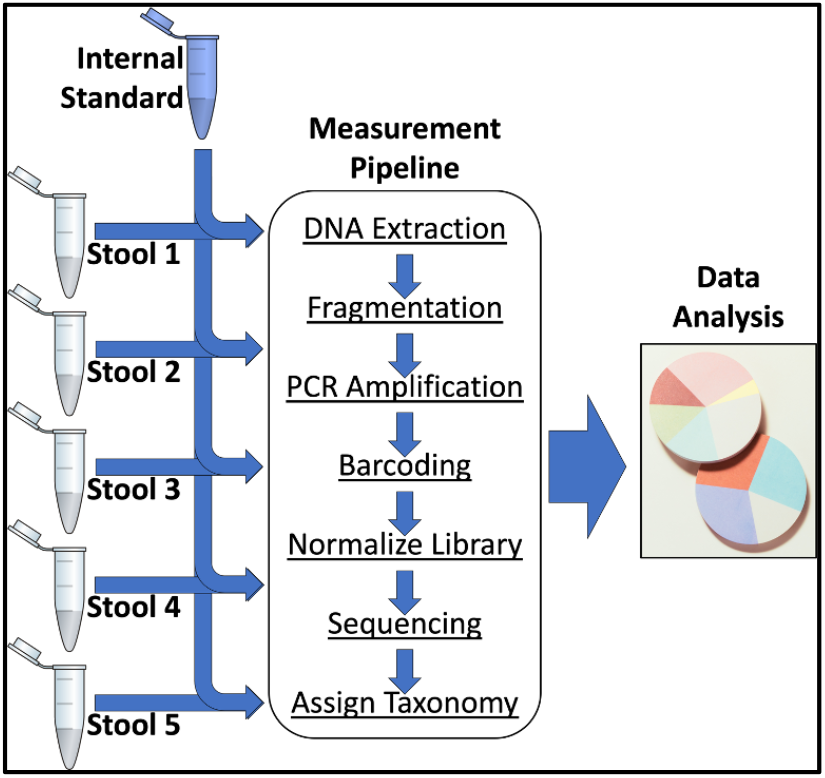
Experimental design specifying systematic addition of an exogenous internal standard microbe into every sample ahead of metagenomic sequencing analysis.

Comparison of measured relative abundances of native taxa to that of the internal standard then corrects for compositionality as shown in Equation 3, but without the ambiguity of referencing two native taxa that may change independently of each other between samples.

Thus, Equation 3 can be rearranged to calculate a ‘Scaled Abundance’ metric as the product of the internal standard actual abundance and the ratio of each native taxa’s relative abundance to the internal standard relative abundance. These Scaled Abundances are then predicted to be directly proportional to each taxa’s actual abundance, with a constant of proportionality that directly reflects the ratio of biases for that taxa and the internal standard:

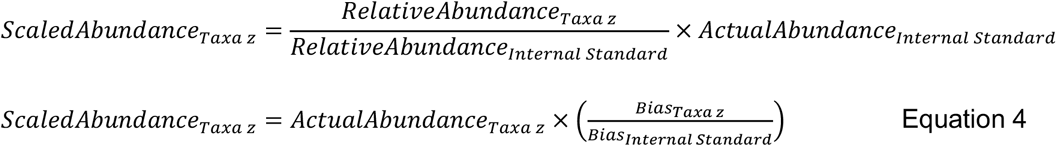

Because biases for each taxa arise from the specifics of the protocol steps employed, the bias ratio in Equation 4 will be constant within locked-down protocols, allowing direct comparisons of abundances between samples, independent of sample compositionality.

There are two primary applications for using this Scaled Abundance metric. In the first scenario, taxa of interest may have been previously identified, isolated, and cultured. In this case, the taxa of interest can be spiked into samples to allow calculation of the bias ratio in Eqn 4. This in turn allows the Scaled Abundance measured in unknown samples to be directly converted into accurately measured actual abundances. However, for many native taxa, bias ratios cannot be independently determined for taxa of interest (e.g., unknown or unculturable taxa). Nevertheless, in this scenario, the biases still remain constant between samples (analyzed using a common protocol), so fold changes in scaled abundance between samples still directly reports fold-changes in actual abundance:

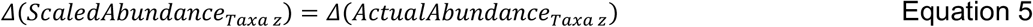

This mathematical framework for using internal standards to calculate a Scaled Abundance metric from MGS data is further explored in the remainder of this manuscript with actual MGS data. With previously published MGS data from a series of microbial mock communities, we reanalyze the results by treating one constituent taxa as an internal standard and showing that the calculated Scaled Abundances correlate well with known actual abundances. Then, we prepared and analyzed 5 distinct fecal samples to demonstrate that the calculated Scaled Abundance is independent of sample composition (as predicted by Equation 4). We further show how to calculate bias ratios for independently characterized taxa, thereby allowing direct measurement of Actual Abundances for known taxa, and how to use fold changes in Scaled Abundance to accurately report fold changes in Actual Abundance for all taxa (as predicted in Equation 5). Finally, we explored a dilution series to show that Scaled Abundance accurately tracks changes in actual abundance for native taxa, even when sample composition is constant (as predicted by Equation 4). In each evaluation, analysis of real data supports the mathematical framework described here and demonstrates that the use of an internal standard to calculate Scaled Abundances accurately corrects for MGS compositionality and enables abundances of individual taxa to be quantified and meaningfully compared between microbiome samples.

### Previously published data

In 2015, Brooks et al. systematically generated a series of 80 combinations of 7 specific bacterial strains commonly associated with the vaginal microbiome.^26^ These diverse metagenomic samples were then analyzed using a common measurement pipeline, including DNA extraction, PCR amplification of the V1-V3 region of the rRNA gene, followed-by next-generation sequencing. While the resulting measured relative abundances exhibited poor concordance with the known actual abundances of taxa, the data was systematically collected, annotated, and made publicly available, and these data have been usefully reevaluated to validate ratiometric approaches to compositional data analysis.^25^ We have reanalyzed this dataset in the current framework by designating one added strain as an internal standard and using its known abundance to calculate scaled abundances for the remaining taxa (Figure 2).

**Figure 2.**
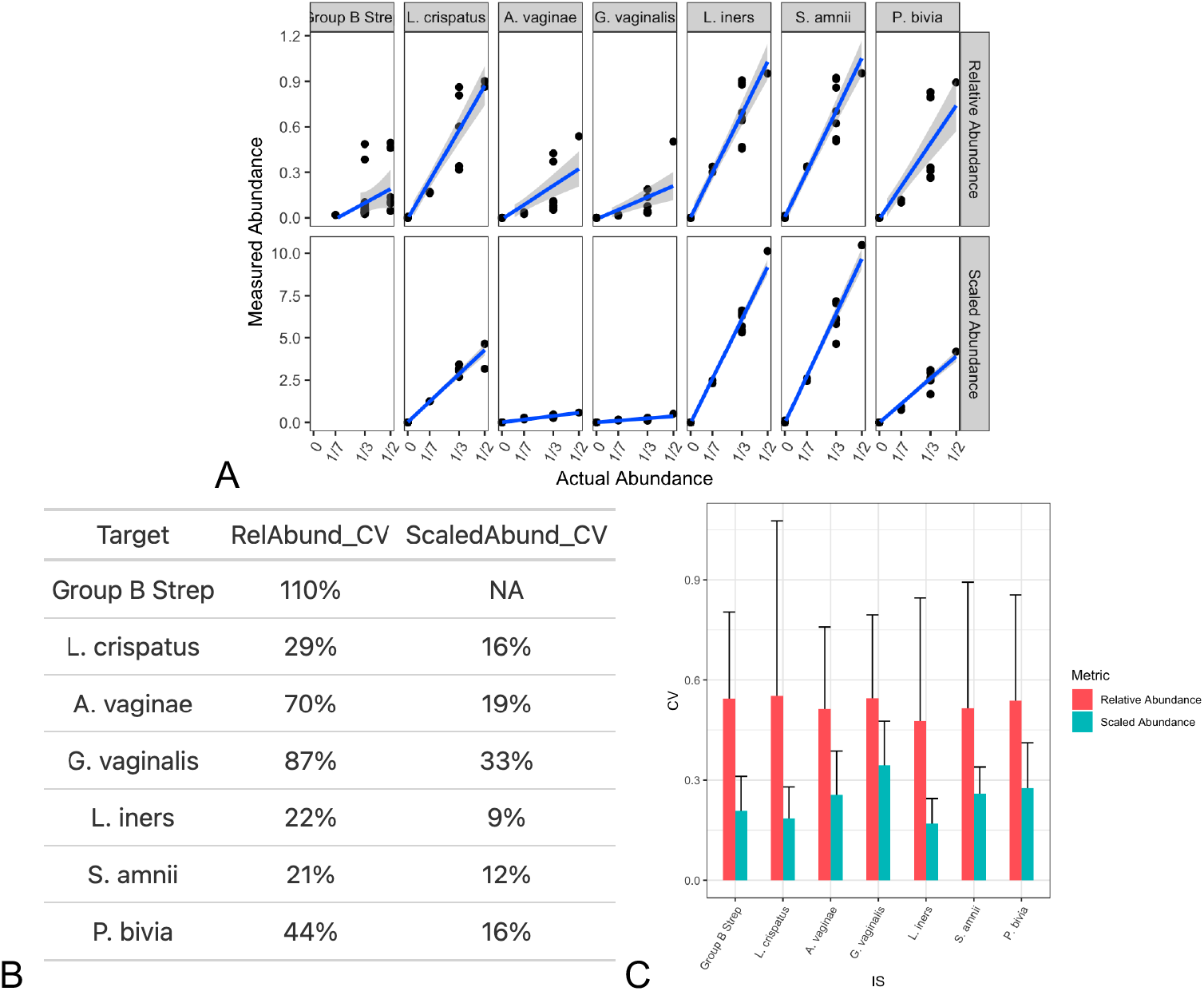
Previously published data from a series of microbial mock communities (Brooks, et al., 2015) was reanalyzed using the mathematical framework described herein by treating one strain as an internal standard for calculating Scaled Abundances. Strain abundances, as represented by relative abundance (RA) or Scaled Abundance (SA), were plotted against their experimentally specified actual abundances (A). (Group B Strep was treated as the internal standard, so no Scaled Abundances were calculated.) Linear regressions and 95% confidence bounds show best fit linear correlation models. The regression goodness-of-fits are summarized (B) as calculated coefficients of variation (CVs) for each regression. This procedure was generalized by systematically treating each strain as the internal standard, and calculating the average goodness-of-fit (and 95% confidence intervals) for remaining strains as measured by each metric (C).

As reported in the analysis from the original manuscript, measured relative abundance of individual taxa exhibited poor correlation with the actual abundances known from sample preparation (Figure 2a, relative abundance). However, when we designated one of the taxa (e.g., Group B Strep) as an internal standard to correct for compositionality, the calculated scaled abundances for the remaining taxa were much better correlated with the known actual abundances (Figure 2a, scaled abundances). Regressions for each taxa’s scaled abundance demonstrated directly proportionality with their actual abundances, as predicted by Equation 4.

The slopes of the scaled abundance regressions varied between taxa, consistent with varying constants of proportionality in each case as predicted by the bias rations in Equation 4. For known, independently quantified taxa (such as the ones explored in this dataset) these regressions provided a quantification of the bias ratio and, in turn, a calibration for the determination of actual abundances from scaled abundances measurements of any future unknown samples. This calibration will be particular to the specific method/protocol employed.

For each taxa, the degree of agreement between the observed and actual abundances (goodness-of-fit) was summarized using a calculated coefficient of variation (CV, Figure 2b). Using Group B Strep as the internal standard, the CVs for relative abundance measurements were quite large and consistently higher than the CVs for Scaled Abundances, presumably because the Scaled Abundance corrected for sample compositionality. Recognizing that any of the mock community taxa could have been treated as the internal standards, these analyses were repeated for each taxa. The average of CVs from the remaining taxa showed that Scaled Abundance, calculated using the internal standards, consistently produced better correlation (lower CVs) between measured and actual abundances than relative abundances (Figure 2c). Taken together, these results were entirely consistent with the described mathematical framework and demonstrated that using internal standards to calculate Scaled Abundances significantly improved MGS quantitation and the ability to accurately correlate individual taxa measurements with known actual abundances.

### Testable Hypothesis 1: Scaled Abundance is independent of sample composition

One of the predictions arising from Equation 4 is that Scaled Abundance measurements for individual taxa will be independent of sample composition. This is in direct comparison to Relative Abundance measurements which are compositionally dependent (as shown in Equation 2). To test this hypothesis, a series of samples were prepared wherein constant amounts of microbial DNA from 4 exogenous taxa were spiked into individual fecal samples exhibiting identical compositions (i.e., technical replicates) or varied compositions (Figure 3). The concentrations of the spiked-in DNA was systematically varied by taxa across 2 orders of magnitude, and it was then uniformly added to each sample. For each sample, MGS analysis yielded Relative Abundance measurements which were used to calculate Scaled Abundances for the spiked-in taxa through comparison to an internal standard as described in Equation 4.

**Figure 3.**
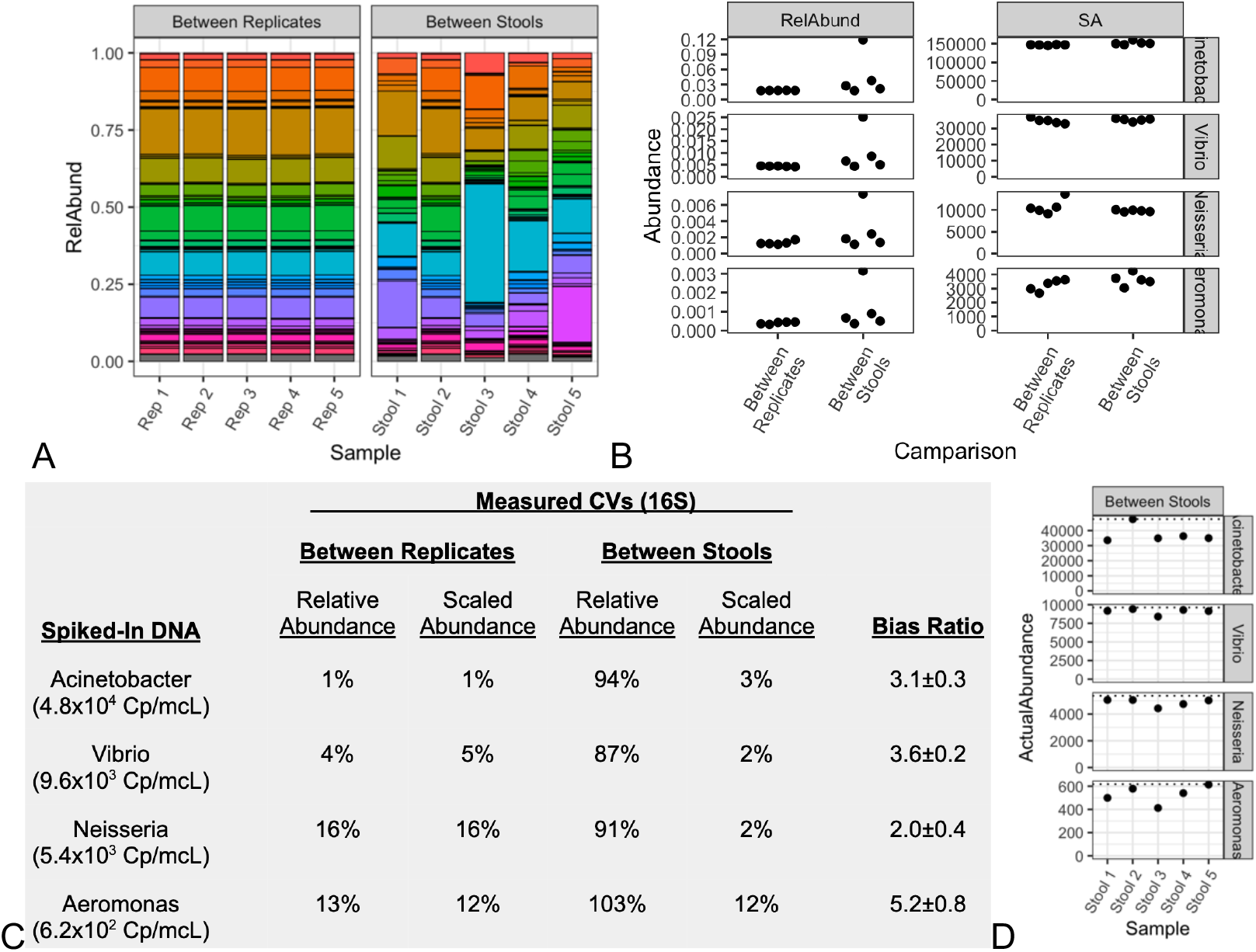
Metagenomic sequencing of diverse stool samples. Stacked barcharts (A) of the most abundant genera reveal high reproducibility within replicate analyses of a single stool sample, and significant compositional differences between stool samples from different donors. Known concentrations of DNA from 4 taxa were uniformly spiked-in, and their individual abundances (B) are plotted for the relative abundance (RelAbund) and Scaled Abundance (SA) metrics and grouped for comparisons between replicates and between distinct stools. The precision of these abundance measurements are tabulated (C) as calculated CVs. Data from the Scaled Abundance technical replicates were used to calculate Bias Ratios (± 95% confidence interval), as described in Equation 4. Using these Bias Ratios, the actual abundances for the ‘between stools’ data are plotted (D) and correlate well with their actual abundances (dotted lines). Shown here for 16S MGS analysis, the same samples were analyzed using a shotgun MGS analysis pipeline with substantially similar results (Figure SI 1).

As expected for MGS analysis of fecal samples, good reproducibility was observed between replicate analyses, while significant differences were observed between five different stool samples (Figure 3A). The inherent compositionality of this measurement is implicit in the way bar chart Relative Abundances always sum to unity. Stacked bar chart plots are particularly useful for observing technical precision or compositional shifts within all of the taxa present.

However, Relative Abundance has limited utility for evaluating individual taxa between samples because of its compositional dependence. Here, when individual spike-in taxa were uniformly added to all samples, their measured Relative Abundances (Figure 3B, RelAbund) were only consistent within a common sample composition. Even though the spiked-in DNA was known to be constant, the observed relative abundances between different fecal samples exhibited significant variation. Further, these variations were strongly correlated between taxa. This behavior is entirely consistent with the expected effect of sample compositionality on the relative abundance measurement as described in Equation 2. In contrast, Scaled Abundances were calculated using an internal standard in an effort to correct for compositionality (Equation 4). Unlike the variation observed for Relative Abundance measurements, the calculated Scaled Abundances exhibited consistency for both technical replicates and across differing sample compositions (Figure 3B, ScaledAbund). This supported the hypothesis that the Scaled Abundance metric would be wholly independent of sample composition.

The precision of observed abundances (by Relative Abundance and Scaled Abundance) were calculated as the coefficient of variation (CV) across each group of samples and for each spiked-in taxa (Figure 3C). Within the measured Relative Abundances of technical replicates, CVs ranged 1%-15%. This precision generally decreased alongside the spike-in concentration, and these were interpreted as representing a lower limit of precision for sample preparation (i.e., spiking in DNA) and handling. When considering the measured Relative Abundances across varied sample compositions, much higher variability (90%-100% CV) was observed for the spiked-in taxa, consistent with the effects of compositionality. In contrast, calculated Scaled Abundances exhibited generally high precision (CV ≤ 15%) for all spiked-in taxa, both within technical replicates and between different sample compositions. Importantly, this showed that the Scaled Abundance approach described herein was able to consistently account for compositionality and achieve high precision across at least 2 orders of magnitude, and even at low relative abundances (RA ∼ 0.0005).

### Accounting for Bias

While the calculated Scaled Abundances succeed at accounting for differences between sample compositions, they are not without bias. Indeed, Equation 4 predicts that the constant of proportionality between the Scaled Abundances for each taxa and their Actual Abundances arises from the ratio of the analytical biases between the specified taxa and the internal standard. Using the known Actual Abundances of each spike-in taxa, bias ratios (*Bias*_*Spike In*_: *Bias*_*Internal Standard*_) were calculated from the technical replicate fecal samples (Figure 3C). While the measured biases are expected to be highly protocol-specific, they were highly reproducible across replicate samples for a locked-down protocol. These bias ratios were then used to calculate actual abundances, from the Scaled Abundances in each distinct stool sample, and exhibited excellent agreement to ground truth (Figure 3D). These results demonstrated that for taxa that can be independently measured and spiked-in, bias ratios can be directly quantified to calibrate the calculated Scaled Abundances to actual abundance determinations.

However, many taxa native to microbiome samples may not be available for independent culture and enumeration. An alternative strategy to account for bias is to recognize that bias is invariant within a specified protocol and will skew Scaled Abundances equally within each sample (Equation 4). Thus, the fold change in Scaled Abundance between samples allows these biases to cancel, thereby directly reporting the fold change in Actual Abundance (Equation 5). When Δ (*Scaled Abundance*) was calculated between the five fecal samples for any of the uniformly added spike-in taxa, the results hovered around unity and were independent of the particular taxa or samples considered (Figure 4A). These results were consistent with an absence of any significant effect of sample composition, taxa identity, or spike-in actual abundance on using internal standards and Scaled Abundances to quantify Δ(*Actual Abundance*) between samples. This result improves on previous ratiometric approaches in that the use of an internal standard reference allowed the comparisons between samples to be directly attributed to individual taxa. The calculated Δ (*Scaled Abundance*) for each of the spike-in taxa as well as the most abundant 10 native taxa are also tabulated (Figure 4B). For native taxa, the fold-changes varied significantly between pairs of samples, as expected for diverse fecal samples.

**Figure 4.**
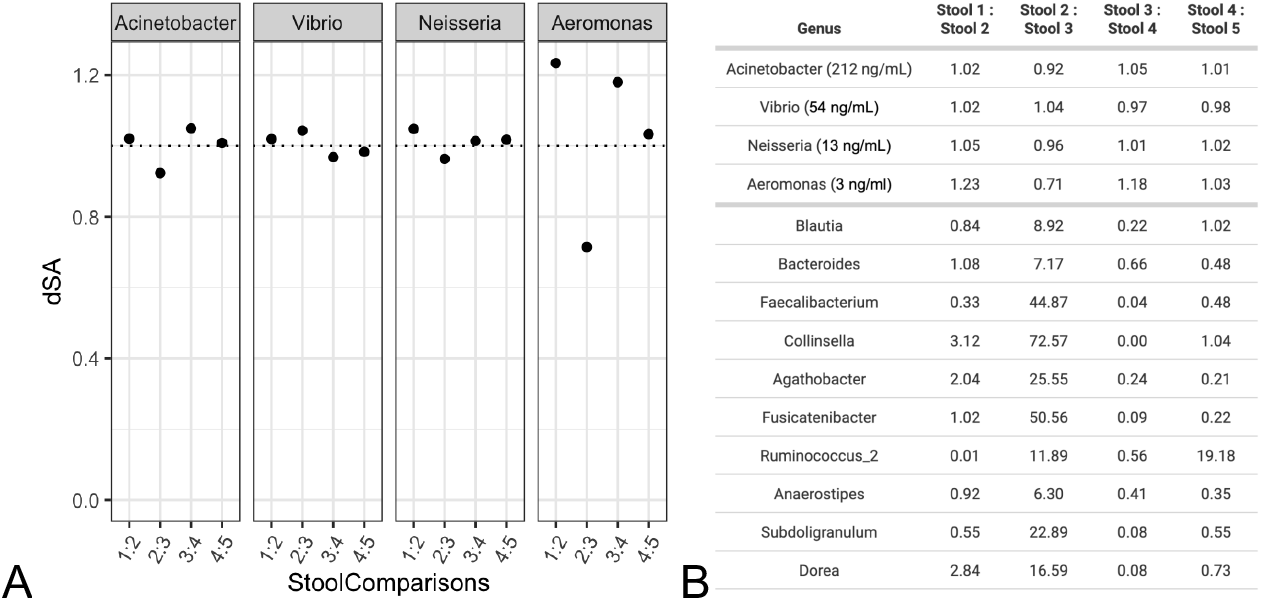
Δ (Scaled Abundance) for each spike-in taxa was calculated and plotted (A) for pairs of the stool samples shown in Figure 3. Since the spike-ins were uniformly added to all samples, Δ (Scaled Abundance) was expected to be unity (dotted line). The table (B) provides calculated Δ (Scaled Abundance) for each of the spike-in taxa and the top 10 most abundant taxa quantified in all 5 stool samples. Shown here for 16S MGS analysis, the same samples were analyzed using a shotgun MGS analysis pipeline with substantially similar results (Figures SI 2 and SI 3). An exhaustive list of all taxa and their measured Δ (Scaled Abundance) values is also available (Table SI 1).

When the same fecal samples were analyzed using shotgun MGS (instead of 16S MGS), substantially similar results were attained (Figure SI 2). In particular, the quantified Δ (*Scaled Abundance*) values for native taxa between samples were nearly identical (Figure SI 3, Table SI 1), suggesting that these results were accurately reporting Δ (*Scaled Abundance*) and reflect the real biological differences in native taxa abundances between samples.

Thus, while bias is unavoidable in MGS measurements, the ratio of the biases between each taxa and the internal standard were consistant between samples and also represented the constant of proportionality between that taxa’s calculated Scaled Abundances and its Actual Abundance (Equation 4). For independently enumerated taxa, this Bias Ratio can be calculated, allowing Scaled Abundances to be accurately converted into measures of Actual Abundance. For other taxa, such as unculturable native species, comparisons between samples still allow accurate quantification of fold changes in actual abundance (Equation 5). In either case, this experimental design for the addition of internal standards and mathematical framework to calculate Scaled Abundances provides a rigorous and quantitative strategy for directly comparing taxa actual abundances between samples.

### Testable Hypothesis 2: Scaled Abundance is proportional to actual abundance

To further demonstrate the utility of an experimental design which includes internal standards, a series of samples were prepared wherein the sample composition was kept constant even while microbial abundances were systematically varied. This was accomplished by systematically diluting a fecal sample into Tris-EDTA buffer (Figure 5a). As can be seen from the stacked bar charts of taxa relative abundances, as measured by shotgun MGS, sample composition remained unchanged across all samples, even while the Actual Abundances of native were known to be decreasing based on sample preparation. This dilution series was then treated as a series of independent samples, into which the internal standard was uniformly added. For these samples, the dilution fraction served as a stand-in for taxa actual abundance when comparing between samples, and measured abundances were correlated with dilution fraction as described in Equation 2 and Equation 4.

**Figure 5.**
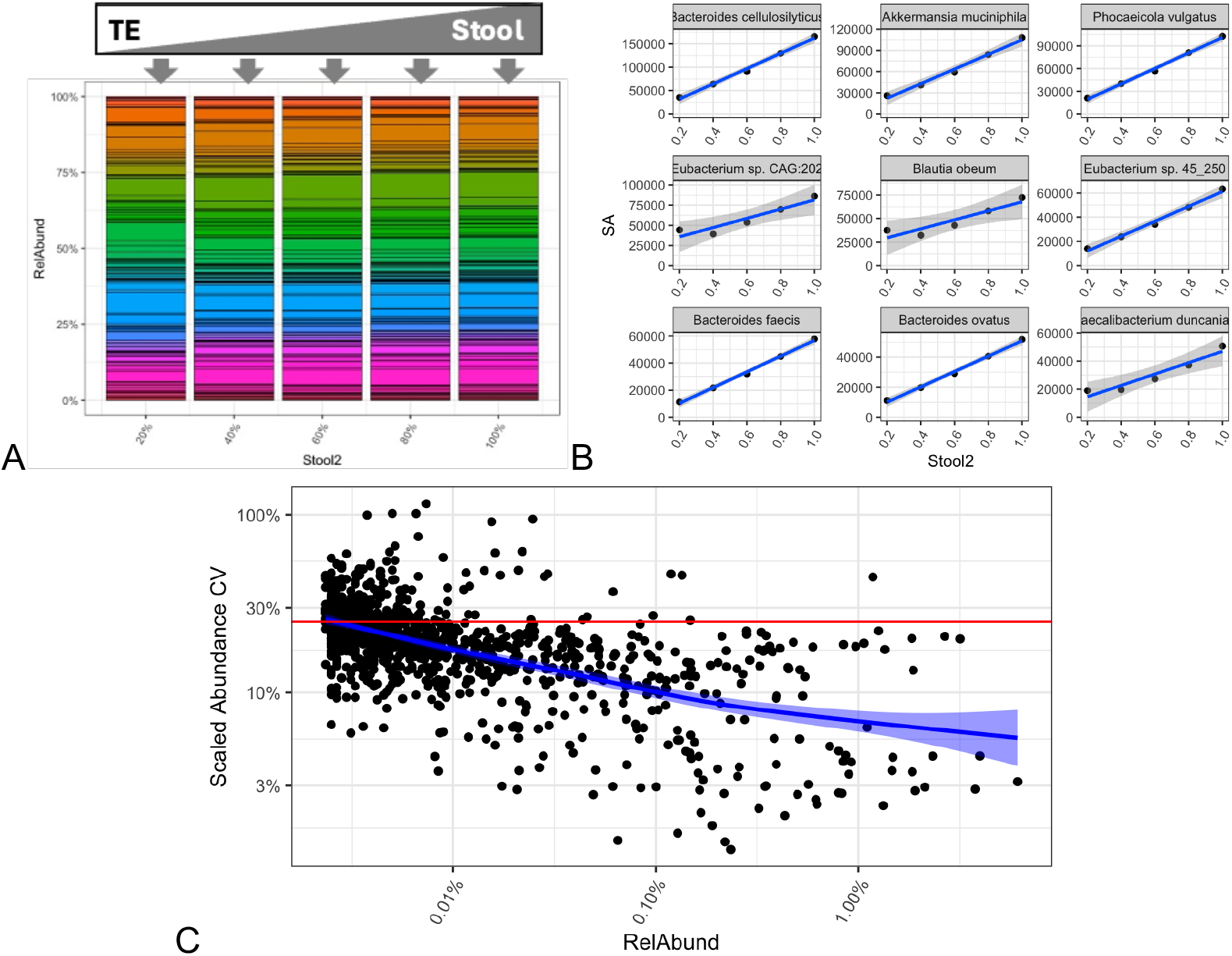
Metagenomic sequencing of a Stool dilution series. Stacked bar charts (A) of a systematic dilution of stool with TE buffer show a consistent sample composition. Comparisons of measured Scaled Abundance to actual stool concentration (B, for the nine most-abundance native taxa) were highly correlated. (Relative abundances for the same taxa exhibited poor correlations with stool concentration, as seen in Figure SI 4.) Similar regressions were determined for all taxa native to the stool, and their goodness-of-fits were plotted (C) as a function of each taxa’s measured relative abundance. Across all taxa (blue trend line), these regressions revealed high correlation (CV≤25%) between measured Scaled Abundances and known actual abundances across a wide range of taxa relative abundances.

When individual Scaled Abundances of the most abundant native taxa were compared to each sample’s known fecal concentration, good correlations were observed, as predicted from Equation 4 (Figure 5B). In contrast, relative abundances of the same taxa exhibited poor correlations with fecal concentration which were generally indistinguishable from the null hypothesis of no correlation (Figure SI 4).

While regressions only from the 10 most abundant taxa are shown, linear fits of Scaled Abundance to fecal concentration were calculated for each native taxa, and a goodness-of-fit was measured for each regression as a coefficient of variation (CV) and plotted against taxa relative abundance in whole stool (Figure 5C). Across a wide range of taxa abundances (down to a relative abundance of 0.00003), the calculated Scaled Abundances were generally (blue trace in Figure 5C) accurate in reporting changes to taxa actual taxa abundances with high precision (CV ≤25%). Taken together, these data show that the calculation of Scaled Abundances, as described in Equation 4, was useful in quantifying the actual abundances of all taxa in the sample, even for low-abundance taxa.

## Conclusions

This study introduces a new experimental design for improving the reliability of microbiome analyses by incorporating internal standards to account for compositional distortions inherent in metagenomic sequencing data. The mathematical framework developed herein demonstrates that the inclusion of an exogenous internal standard, whose actual abundance is measurable, allows for the calculation of taxa-specific ‘Scaled Abundances’ that are (i) independent of sample composition and (ii) directly proportional to actual biological abundances. This approach is consistent with prior ratiometric analysis strategies, with the added simplicity of accounting for compositionality through comparison to a single, well-characterized taxa systematically added at a known actual abundance.

For example, while conventional relative abundances can be used to detect compositional shifts among all taxa in aggregate and to categorize whole microbiome samples (e.g., case-control comparisons), the calculated Scaled Abundance described herein will enable researchers to evaluate individual constituent taxa and quantify changes in taxa actual abundances between samples. This ability to quantitatively interrogate individual taxa is central for hypothesis generation and understanding how microbial composition relates to observed microbiome function. In longitudinal studies with locked-down protocols, Scaled Abundances will enable accurate tracking of individual microbial abundance over time (e.g., following a defined intervention). Finally, by correcting for MGS compositionality, the framework described here opens new opportunities for quantitatively characterizing conserved sub-microbiomes found within larger, variable microbiome compositions.^42^

Through rigorous analysis of previously published mock community data and freshly analyzed gut microbiome samples, we show that calculated Scaled Abundances outperform traditional relative abundance measurements in both precision and accuracy, and demonstrate their potential for quantitative metagenomic profiling. Specifically, Scaled Abundances calculated for MGS characterization of mock communities agreed much better than raw relative abundances with taxa actual abundances across varied compositions. When exogenous taxa were spiked into fecal samples at known actual abundances spanning several orders of magnitude, the Scaled Abundance metric accurately reflected actual abundances and was wholly independent of sample composition. Furthermore, the use of internal standards allowed accurate comparisons of native taxa between samples, even when taxa-specific analytical biases could not be independently measured. Finally, high precision and accuracy was demonstrated by using Scaled Abundances to track actual abundances of native taxa across a series of stool dilutions, even for low-relative abundance taxa.

The results presented here validate the proposed experimental design and mathematical framework that uses routine, systematic addition of internal standards to microbiome samples to correct for compositionality in MGS analyses. This is particularly crucial in microbiome studies where variability in sample composition often complicates the interpretation of data. This approach is flexible and is shown to be applicable to both shotgun and amplicon-based sequencing methods, which further broadens its utility across a wide range of microbiome research areas. Moreover, by offering a pathway to accurate and reproducible abundance measurements at the level of individual taxa, this methodology could play a pivotal role in advancing microbiome research, particularly in clinical and environmental health contexts where precise taxa quantification is essential for modeling diverse biological phenomena.

## Methods

### Brooks, et al. (2015) data

Amplicon based MGS analyses of 80 distinct mock communities of 7 microbial taxa were downloaded from the original publication.^26^ The downloaded results provide relative abundances for each taxa in each mock community. These data were reanalyzed in R (version 4.3.2) here by treating *‘Group B Strep*’ as an internal standard to allow calculation of Scaled Abundances for the remaining taxa. (Mock communities that didn’t include *Group B Strep* were omitted from further analysis.) The specified dilutions from overnight stock (e.g., ½, ⅓, 1/7) were taken as the known actual abundances for each taxa and correlated to measured relative abundances or Scaled Abundances. The goodness-of-fit of these regressions were summarized as coefficients of variation (CVs) for each taxa. This procedure was repeated treating alternate strains as the ‘internal standard’, and the average of CVs in each case was used to compare goodness-of-fits with respect to internal standard selection. The raw data and full code used for this analysis has been made publicly available (doi.org/10.18434/mds2-3760).

### Human microbiome sample prep

The human microbiome samples used in this study were used previously in the Mosaic Standards Challenge and have been described previously.^29,30^ In short, 5 fecal samples were collected from 5 different anonymized donors. Each sample was prepared by polling and homogenizing multiple bowel movements from each donor, stabilizing the mixtures with Omnigene Gut Solution, and preparing 1 mL identical aliquots at a final concentration of 100 mg/mL. Fecal aliquots were stored at -80 C until ready for analysis. Dilution series were generated by mixing the thawed samples into Tris-EDTA buffer.

### Internal Standard

The experimental design described here specified the systematic addition of an internal standard prior to the traditional metagenomic sequencing measurement pipeline. We selected DNA from *Legionella pneumophila* as an internal standard due to its absence in previous metagenomic characterizations of the stool samples. Additionally, this material was stable and had been well quantified by ddPCR previously.^43^ Genomic DNA from *Legionella Pneumophila* was added uniformly to all samples for a final concentration of (2.16×10^5^ ± 3.7×10^3^) copies/mcl.

### Spike-in DNA

For experiments where specified concentrations of microbial DNA were spiked-in to fecal samples (i.e., Testable Hypothesis #1), pure genomic microbial DNA was added from *Acinetobacter baumanii, Vibrio furnissii, Neisseria meningitidis, and Aeromonas hydrophila*. These DNA genomes were sourced from NIST RM8376^43^ and were added to fecal samples at the final concentrations of 4.8×10^4^ copies/mcL, 9.6×10^3^ copies/mcL, 5.4×10^3^ copies/mcL, and 6.2×10^2^ copies/mcL, respectively.

### Metagenomic sequencing measurement pipeline

#### DNA extraction

DNA was extracted using the ZR Fecal DNA miniprep (cat# D6010) following the manufacturer protocol. Briefly, each sample was combined with 400mcL of the lysis buffer and vortexed (MoBio Genie 2) for 20 minutes at full speed. Lysis tubes were centrifuged at 10,000xg for 1 minute, and the lysate was processed through the spin filter at 7,000xg for 1 minute. The filtrate was added to a 1.2 mL DNA binding buffer, mixed by pipetting, and centrifuged in batches at 10,000xg through the spin column. The bound DNA was washed and eluted into 150 mcL of elution buffer. The extracted DNA was quantified by fluorescence using the DeNovix DS-11FX fluorometer with the DeNovix dsDNA High Sensitivity kit (Catalog #: KIT-DADNA-HIGH-2).

#### Library Preparation

Next-generation sequencing libraries were prepared for both shotgun and 16S amplicon analyses. For shotgun sequencing, extracted DNA was fragmented, amplified, and barcoded using the Nextera XT DNA Library Preparation Kit (Illumina) and Nextera XT Index kit V2 (Illumina, catalog # 15052163) as specified by manufacturer protocols. For amplicon sequencing, the 16S rRNA gene was amplified using primers for the V4 variable region sourced from IDT (10mcM RxnRead Primer Pool). 1mcL of the primer pool was combined with 12.5 mcL of Kapa HiFi HotStart readymix (KapaBiosystems, Catalog # 07958935001) and 12ng extracted DNA in a final volume of 25mcL for PCR (initial denaturation (180s at 95 °C), 18 cycles of denaturing (30s at 98 °C), annealing (15s at 55 °C), and elongation (20s at 72.0 °C), and a final extension (300s at 72.0 °C)). The 16S amplicons were purified using SPRIselect beads (Beckman Coulter, Catalog # B23318) at a 0.8:1 ratio of Beads:amplicon, washing beads with molecular grade ethanol and resuspended in pure water. Finally, barcodes were added (Nextera XT Index kit V2) as specified by manufacturer protocols. For both shotgun and amplicon library prep, samples were quantified by fluorescence (DeNovix) for DNA yield, and 10 ng from each sample were pooled for sequencing.

#### Sequencing

Pooled libraries were quantified by fluorescence, diluted to 4nM, and denatured following manufacturer protocol (Document # 15039740 v10, Protocol A). Denatured libraries were further diluted to 12 pM, combined with a 5% PhiX (V3 cat# 15017666 from Illumina), and sequenced by paired-end sequencing on an Illumina MiSeq (MiSeq Reagent Kit v3 600-cycle, cat #: MS-102-3003).

### Initial Bioinformatic analysis

Demultiplexing and adapter trimming was completed as part of the Illumina MiSeq Generate FASTQ workflow. Fastq files were analyzed separately for shotgun and amplicon sequencing, as described subsequently. The final output of both shotgun and amplicon bioinformatic analysis was a table specifying the taxa observed in each sample and their measured relative abundances.

#### For shotgun sequencing

BBduk (38.90)^44^ was used to quality filter the raw data, and paired-end sample data were analyzed using Centrifuge^45^ with the Web of Life database^46^. The bash script files have been made publicly available (doi.org/10.18434/mds2-3760).

#### For amplicon sequencing

Cutadapt (2.8)^47^ was used to remove primer sequences, and DADA2 (1.20.0)^48^ was used to account for sequencing errors, determine absolute sequence variants, measure relative abundances, and assign taxonomy based on the Silva database (version 132)^49^. The raw data and code (R, version 4.3.2)^50^ used for amplicon sequencing analysis has been made publicly available (doi.org/10.18434/mds2-3760).

### Scaled Abundance calculations

The definitions and mathematical framework for calculating the Scaled Abundance metric from tables of identified taxa from each sample and their measured relative abundances is provided in the results section of this manuscript. These operations were implemented in R (version 4.3.2), and all raw data and the code have been made publicly available (doi.org/10.18434/mds2-3760).

## Data availability

All raw data and analysis code described in this manuscript has been made publicly available (doi.org/10.18434/mds2-3760).

## Disclosures and acknowledgements

All work was reviewed and approved by the U.S. National Institute of Standards and Technology (NIST) Research Protections Office. This study (protocol # MML-2019-0135) was determined to be “not human subjects research” as defined in the Common Rule (45 CFR 46, Subpart A).

This effort received no funding from commercial or not-for-profit sectors and was supported entirely by NIST. Certain commercial equipment, instruments, and materials are identified in this manuscript to foster understanding. Such identification does not imply recommendation or endorsement by NIST, nor does it imply that the materials or equipment identified are necessarily the best available for the purpose.

## Author contributions

SPF conceptualized the project with input from SLS and JGK. Experiments were designed by SPF and SLS and executed by MEH and JND. Initial bioinformatic analysis was done by JGK and SLS, and Scaled Abundances calculations and final bioinformatic analyses were done by SPF. SPF wrote the manuscript and prepared the figures. In writing this manuscript, ChatGPT was used to generate initial drafts of the conclusions and abstract sections that were then edited extensively by the authors. All authors read and approved the final manuscript.

