## Supplementary material for "A mathematical framework to correct for compositionality in microbiome datasets": SI material: Figure SI 1, Figure SI 2, Table SI 1, Figure SI 3, and Figure SI 4.

**SI Figures and Tables included:**

- ***Figure SI 1:*** *Metagenomic sequencing of diverse stool samples, by shotgun MGS*
- ***Figure SI 2:*** $\Delta(ScaledAbundance)$*, by shotgun MGS*
- ***Table SI 1:*** *Exhaustive summary of* $\Delta(ScaledAbundance)$ *for all taxa*
- ***Figure SI 3:*** Comparison of $\Delta(ScaledAbundance)$ *for shotgun and amplicon MGS*
- ***Figure SI 4:*** *Comparisons of measured relative abundance to actual stool concentration.*

***Figure SI 1:*** *Metagenomic sequencing of diverse stool samples: this figure shows analysis of the same samples as Figure 3 from the main manuscript, but with shotgun MGS analysis (instead of amplicon MGS). Stacked barcharts (A) of the most abundant genera reveal high reproducibility within replicate analyses of a single sample, and significant compositional differences between stool samples from different donors. The individual abundances of spiked-in microbial taxa (B) are plotted for the relative abundance and Scaled Abundance metrics and grouped for comparisons between replicates and between stool samples. The precision of these abundances are tabulated (C) as a calculated coefficients of variation (CVs) alongside the bias ratio calculated from the 'between replicates' data based on equation 4 and the known spike-in concentrations. Using these Bias Ratios, the actual abundances for the 'between stools' data are calculated (D) and correlate well with the expected abundance (dotted lines).*

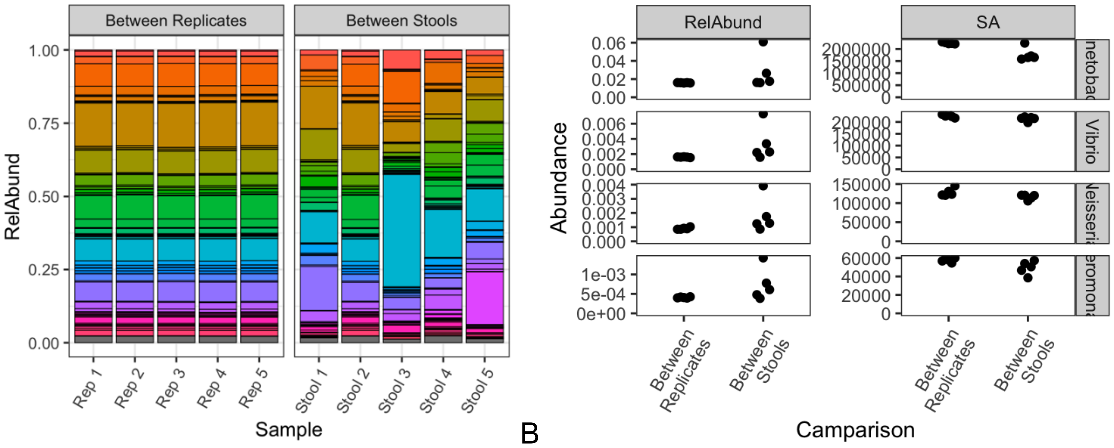


A

| Spiked-In DNA | Measured CVs (WGS) | | | | |
|---|---|---|---|---|---|
| | Between Replicates | | Between Stools | | |
| | Relative Abundance | Scaled Abundance | Relative Abundance | Scaled Abundance | Bias Ratio |
| Acinetobacter (212 ng/mL) | 1% | 2% | 70% | 15% | 10613±204 |
| Vibrio (54 ng/mL) | 3% | 3% | 69% | 4% | 4210±164 |
| Neisseria (13 ng/mL) | 8% | 8% | 57% | 6% | 9661±963 |
| Aeromonas (3 ng/ml) | 3% | 4% | 56% | 15% | 17356±819 |

C

D

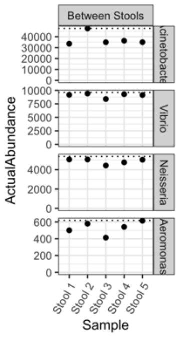

***Figure SI 2:* Δ*(ScaledAbundance), by shotgun MGS. This figure shows analysis of the same samples as Figure 4 from the main manuscript, but with shotgun MGS analysis (instead of amplicon MGS). $\Delta(ScaledAbundance)$ between pairs of stool samples was plotted (A) for each spike-in and agreed well with $\Delta(ActualAbundance)$. (Since the spike-ins were uniformly added to all samples, $\Delta(ScaledAbundance)$ should be unity in each comparison.) The table (B) provides calculated $\Delta(ScaledAbundance)$ for each of the spike-in taxa and the top 20 most abundant taxa quantified in all 5 stool samples.*

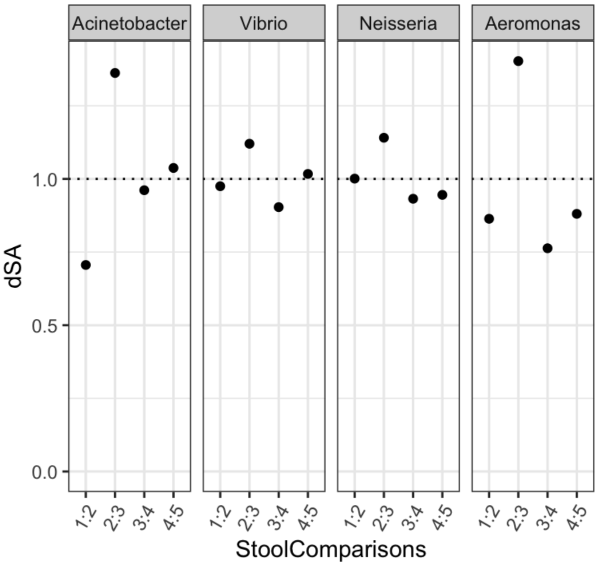


A

B

| Genus | Stool 1 : Stool 2 | Stool 2 : Stool 3 | Stool 3 : Stool 4 | Stool 4 : Stool 5 |
|---|---|---|---|---|
| Acinetobacter (212 ng/mL) | 0.71 | 1.36 | 0.96 | 1.04 |
| Vibrio (54 ng/mL) | 0.98 | 1.12 | 0.90 | 1.02 |
| Neisseria (13 ng/mL) | 1.00 | 1.14 | 0.93 | 0.95 |
| Aeromonas (3 ng/ml) | 0.86 | 1.40 | 0.76 | 0.88 |
| Bacteroides | 0.72 | 10.52 | 0.39 | 0.90 |
| Blautia | 0.94 | 13.51 | 0.17 | 0.70 |
| Phocaeicola | 1.68 | 7.91 | 0.51 | 0.40 |
| Segatella | 1.37 | 3.12 | 0.02 | 0.05 |
| Akkermansia | 0.04 | 3.48 | 126.42 | 0.55 |
| Eubacterium | 0.24 | 54.22 | 0.16 | 0.37 |
| Alistipes | 0.47 | 5.18 | 0.16 | 3.53 |
| Clostridium | 0.24 | 15.37 | 0.15 | 0.62 |
| Faecalibacterium | 0.40 | 26.30 | 0.06 | 0.76 |
| Agathobacter | 1.45 | 37.68 | 0.14 | 0.24 |
| Roseburia | 1.16 | 9.33 | 0.27 | 0.67 |
| Ruminococcus | 0.25 | 1.87 | 0.22 | 4.56 |
| Mediterraneibacter | 1.59 | 11.35 | 0.11 | 0.79 |
| Collinsella | 2.76 | 105.73 | 0.00 | 0.89 |
| Parabacteroides | 0.99 | 14.43 | 0.26 | 0.78 |
| unclassified Bacillota genus | 0.41 | 4.08 | 0.76 | 0.74 |
| Dorea | 2.31 | 11.07 | 0.12 | 0.85 |
| Fusicatenibacter | 1.08 | 31.52 | 0.10 | 0.38 |
| Anaerostipes | 0.91 | 6.23 | 0.36 | 0.54 |
| Anaerobutyricum | 2.56 | 1.39 | 1.72 | 0.20 |

***Table SI 1***: *Exhaustive summary of $\Delta(ScaledAbundance)$ measured between 5 stool samples for all taxa called by both amplicon (16S) and shotgun (WGS) analyses.*

| Genus | MGS Type | Stool Comparison | | | |
|---|---|---|---|---|---|
| | | **1:2** | **2:3** | **3:4** | **4:5** |
| Acinetobacter | 16S | 1.02 | 0.92 | 1.05 | 1.01 |
| Acinetobacter | WGS | 0.71 | 1.36 | 0.96 | 1.04 |
| Vibrio | 16S | 1.02 | 1.04 | 0.97 | 0.98 |
| Vibrio | WGS | 0.98 | 1.12 | 0.90 | 1.02 |
| Neisseria | 16S | 1.05 | 0.96 | 1.01 | 1.02 |
| Neisseria | WGS | 1.00 | 1.14 | 0.93 | 0.95 |
| Aeromonas | 16S | 1.23 | 0.71 | 1.18 | 1.03 |
| Aeromonas | WGS | 0.86 | 1.40 | 0.76 | 0.88 |
| Blautia | 16S | 0.84 | 8.92 | 0.22 | 1.02 |
| Blautia | WGS | 0.94 | 13.51 | 0.17 | 0.70 |
| Bacteroides | 16S | 1.08 | 7.17 | 0.66 | 0.48 |
| Bacteroides | WGS | 0.72 | 10.52 | 0.39 | 0.90 |
| Faecalibacterium | 16S | 0.33 | 44.87 | 0.04 | 0.48 |
| Faecalibacterium | WGS | 0.40 | 26.30 | 0.06 | 0.76 |
| Collinsella | 16S | 3.12 | 72.57 | 0.00 | 1.04 |
| Collinsella | WGS | 2.76 | 105.73 | 0.00 | 0.89 |
| Agathobacter | 16S | 2.04 | 25.55 | 0.24 | 0.21 |
| Agathobacter | WGS | 1.45 | 37.68 | 0.14 | 0.24 |
| Fusicatenibacter | 16S | 1.02 | 50.56 | 0.09 | 0.22 |
| Fusicatenibacter | WGS | 1.08 | 31.52 | 0.10 | 0.38 |

| | | | | | |
|---|---|---|---|---|---|
| Anaerostipes | 16S | 0.92 | 6.30 | 0.41 | 0.35 |
| Anaerostipes | WGS | 0.91 | 6.23 | 0.36 | 0.54 |
| Subdoligranulum | 16S | 0.55 | 22.89 | 0.08 | 0.55 |
| Subdoligranulum | WGS | 0.31 | 23.62 | 0.13 | 0.62 |
| Dorea | 16S | 2.84 | 16.59 | 0.08 | 0.73 |
| Dorea | WGS | 2.31 | 11.07 | 0.12 | 0.85 |
| Akkermansia | 16S | 0.08 | 4.10 | 198.87 | 0.35 |
| Akkermansia | WGS | 0.04 | 3.48 | 126.42 | 0.55 |
| Lachnoclostridium | 16S | 7.08 | 1.24 | 1.63 | 0.29 |
| Lachnoclostridium | WGS | 0.58 | 5.35 | 0.35 | 1.22 |
| Alistipes | 16S | 0.29 | 4.64 | 0.24 | 2.86 |
| Alistipes | WGS | 0.47 | 5.18 | 0.16 | 3.53 |
| Serratia | 16S | 0.31 | 0.07 | 5.66 | 1.91 |
| Serratia | WGS | 0.19 | 13.26 | 0.30 | 0.86 |
| Parabacteroides | 16S | 1.00 | 11.99 | 0.33 | 0.57 |
| Parabacteroides | WGS | 0.99 | 14.43 | 0.26 | 0.78 |
| Streptococcus | 16S | 10.07 | 0.91 | 0.13 | 2.46 |
| Streptococcus | WGS | 3.04 | 1.66 | 0.12 | 2.43 |
| Marvinbryantia | 16S | 0.06 | 9.43 | 0.18 | 2.30 |
| Marvinbryantia | WGS | 0.57 | 1.90 | 0.90 | 1.48 |
| Lachnospira | 16S | 0.83 | 6.37 | 0.93 | 0.24 |
| Lachnospira | WGS | 0.34 | 18.40 | 0.25 | 0.39 |
| Roseburia | 16S | 0.21 | 11.41 | 0.13 | 0.85 |
| Roseburia | WGS | 1.16 | 9.33 | 0.27 | 0.67 |

| | | | | | |
|---|---|---|---|---|---|
| Butyricicoccus | 16S | 1.13 | 42.28 | 0.07 | 0.80 |
| Butyricicoccus | WGS | 0.87 | 22.17 | 0.11 | 0.90 |
| Intestinibacter | 16S | 2.03 | 0.03 | 3.14 | 0.13 |
| Intestinibacter | WGS | 1.82 | 0.34 | 2.34 | 0.16 |
| Phascolarctobacterium | 16S | 1.04 | 123.04 | 5.73 | 0.30 |
| Phascolarctobacterium | WGS | 0.90 | 22.18 | 0.35 | 2.61 |
| Adlercreutzia | 16S | 0.53 | 2.21 | 0.70 | 4.89 |
| Adlercreutzia | WGS | 0.40 | 2.74 | 0.68 | 2.57 |
| Aliivibrio | 16S | 1.09 | 1.38 | 1.04 | 0.85 |
| Aliivibrio | WGS | 1.03 | 1.35 | 1.04 | 0.80 |
| Intestinimonas | 16S | 0.07 | 6.39 | 0.17 | 4.57 |
| Intestinimonas | WGS | 0.17 | 7.28 | 0.18 | 1.93 |
| Agrococcus | 16S | 0.99 | 1.34 | 1.05 | 0.90 |
| Agrococcus | WGS | 0.54 | 5.03 | 0.56 | 1.21 |
| Methanobrevibacter | 16S | 0.00 | 337.73 | 5.84 | 0.81 |
| Methanobrevibacter | WGS | 0.00 | 362.61 | 0.39 | 1.23 |
| Oscillibacter | 16S | 0.18 | 17.05 | 0.10 | 3.79 |
| Oscillibacter | WGS | 0.24 | 11.01 | 0.13 | 1.96 |
| Hespellia | 16S | 1.43 | 3.54 | 0.54 | 1.54 |
| Hespellia | WGS | 0.61 | 11.34 | 0.17 | 0.52 |
| Flavonifractor | 16S | 0.57 | 9.91 | 0.07 | 0.48 |
| Flavonifractor | WGS | 0.65 | 5.73 | 0.10 | 0.56 |
| Turicibacter | 16S | 2.89 | 0.04 | 5.99 | 4.50 |

| | | | | | |
|---|---|---|---|---|---|
| Turicibacter | WGS | 0.91 | 0.43 | 2.55 | 1.58 |
| Hungatella | 16S | 1.19 | 0.52 | 15.51 | 0.17 |
| Hungatella | WGS | 0.63 | 4.01 | 0.50 | 0.91 |
| Actinomyces | 16S | 5.57 | 1.33 | 0.09 | 4.99 |
| Actinomyces | WGS | 0.85 | 4.03 | 0.27 | 1.34 |
| Holdemania | 16S | 0.29 | 19.32 | 0.25 | 0.59 |
| Holdemania | WGS | 0.35 | 14.60 | 0.14 | 1.24 |
| Dielma | 16S | 0.07 | 24.73 | 0.06 | 1.48 |
| Dielma | WGS | 1.17 | 1.48 | 0.88 | 0.35 |

***Figure SI 3:*** *$\Delta(ScaledAbundance)$ between stool samples was determined independently for shotgun and amplicon MGS bioinformatics analyses and were strongly correlated. This supported the prediction of Equation 5 of the main text that $\Delta(ScaledAbundance)$ can be used to accurately quantify fold changes in taxa actual abundances, independent of analysis methodology.*

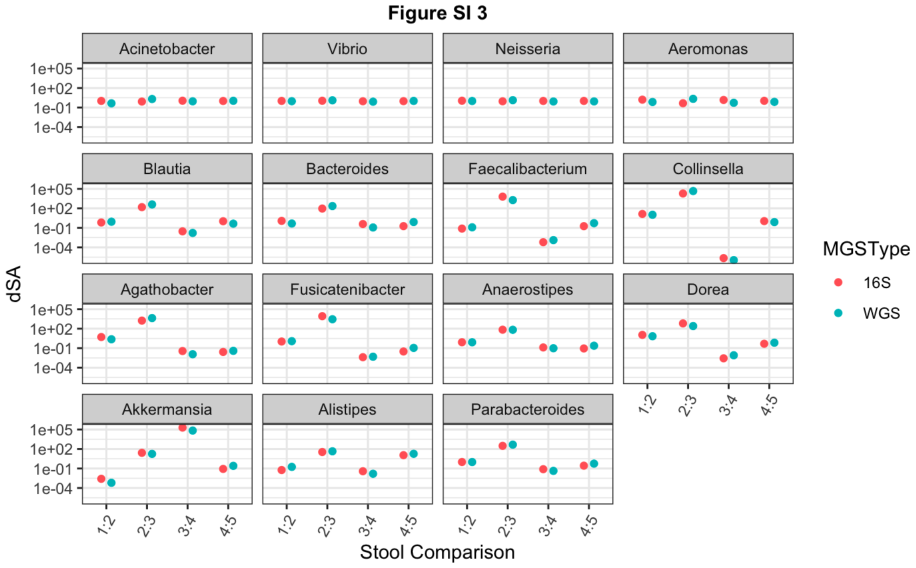

***Figure SI 4:*** *Comparisons of measured relative abundance to actual stool concentration. As a corollary to Figure 5B from the main document, the observed relative abundances were plotted against stool dilution for each of the highest relative abundance genera native to the stool. Linear regressions for each genera exhibited correlation.*

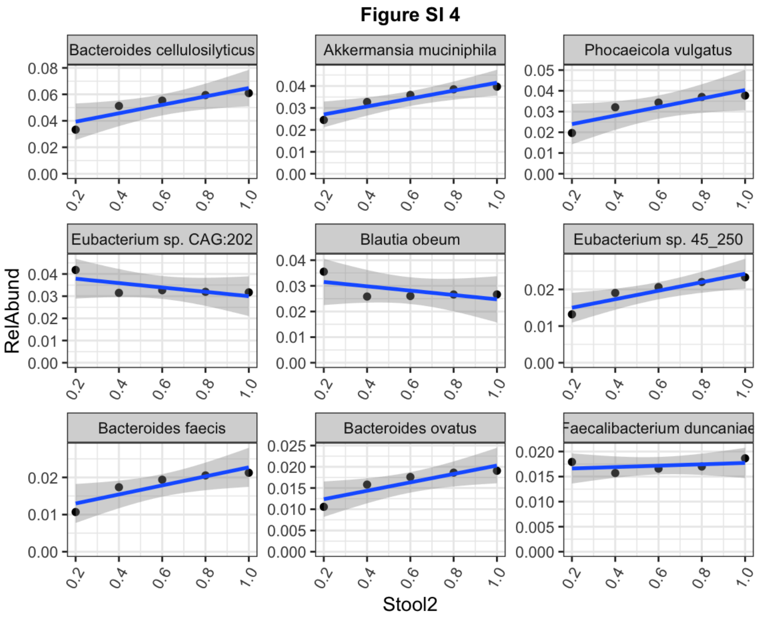
